# Accurate somatic variant detection from formalin fixed, paraffin embedded tissue (FFPE) derived WES and WGS by DeepOmicsFFPE-PLUS, a sequence context-based transformer

**DOI:** 10.64898/2026.02.05.702695

**Authors:** Wan Seop Kim, Ahwon Lee, Inyoung Kim, Seong-Eui Hong, Juyeong Park, Jongsung Lim, Eonji Noh, Minji Kim, Jiin Park, Jiwon Han, Misun Park, Heeyeon Cho, Jonghwan Shin, Yeongyu Yang, Soonmyung Paik, Dong-hyuk Heo

**Affiliations:** Department of Pathology, School of Medicine, Konkuk University, Seoul, Republic of Korea; Department of Hospital Pathology, Seoul St. Mary’s Hospital, College of Medicine, The Catholic University of Korea, Seoul, Republic of Korea; Theragen Bio Co., Ltd., Seongnam-si, Gyeonggi-do, Republic of Korea; Dream Hall, Seongnam Campus, Korea Polytechnic College, Seongnam-si, Gyeonggi-do, Republic of Korea; Severance Biomedical Science Institute, Yonsei University College of Medicine, Seoul, Republic of Korea

## Abstract

Formalin-fixed paraffin-embedded (FFPE) tissues represent a vast archival resource for genomic studies, yet their utility remains constrained by fixation-induced DNA damage and subsequent sequencing artifacts. To comprehensively characterize and address this challenge, we analyzed matched FFPE and fresh-frozen tumor samples from two institutions, spanning different storage durations, DNA qualities, sequencing platforms (WES and WGS), exome capture kits, and somatic variant callers. We found that FFPE-induced artifacts exhibit strong batch- and age-specific patterns, with a predominance of C:G>T:A substitutions, which particularly complicate the accurate identification of low allele frequency true variants. Enzymatic repair methods partially alleviated these artifacts but remained insufficient. To overcome these limitations, we developed DeepOmicsFFPE-PLUS(https://github.com/Theragen-Bio/DeepOmicsFFPE-PLUS), an advanced AI-based tool to accurately distinguish true somatic variants from FFPE-specific artifacts. DeepOmicsFFPE-PLUS demonstrated consistently superior performance across diverse conditions, achieving high sensitivity and specificity—even for low-frequency variants—outperforming existing tools. Application of our model to WGS data further enabled recovery of biologically relevant mutational signatures, including restoration of microsatellite instability (MSI)-associated signatures initially obscured by FFPE artifacts. Our findings underscore the necessity of artifact-aware variant calling in FFPE genomics and establish DeepOmicsFFPE-PLUS as a robust tool for artifact removal, enabling high-fidelity downstream analyses and personalized therapeutic target discovery.

## Introduction

Formalin-Fixed, Paraffin-Embedded (FFPE) samples are widely utilized in histopathological studies and have also become an essential resource for genomic research^1,2^. Due to their widespread availability and the ability to retrospectively analyze patient samples, FFPE-derived DNA is frequently used in next-generation sequencing (NGS) studies for both clinical and research purposes^2^. However, the yield of DNA from FFPE samples is typically low and the NGS data often exhibit uneven coverage across the genome. Specifically, suboptimal reverse crosslinking between DNA and proteins could cause low yield of DNA. Hydrolysis of phosphodiester bonds in the backbone of DNA causes fragmentation making sequence coverage low. The most significant problem is that the fixation and embedding processes, as well as DNA extraction, introduce artifactual modifications, which can lead to incorrect variant identification and impede downstream applications^3^.

Numerous studies have recognized the issue arises in NGS data from FFPE samples and have proposed various experimental and computational approaches to mitigate the problems. A method such as HiTE (Highly concentrated Tris-mediated DNA extraction), employed high-concentration tris buffer which protects DNA damage while improving reverse-crosslinking efficiency significantly improved DNA yield and quality compared to the traditional method^4^. Although this method led to improved genome coverage, the study did not assess the presence or reduction of artifactual alterations commonly observed in FFPE-derived sequencing data. The most frequent artifactual modification is the deamination of cytosine, which results in uracil, subsequently leading to artifactual C:G>T:A mutations^5–7^. An experimental approach has been proposed to repair the cytosine deamination by treating the DNA with uracil-DNA glycosylase (UDG) which generates apurinic/apyrimidinic site (AP site)^8^. The DNA template molecules containing the AP site are excluded from library preparation because DNA polymerase can not synthesize the complementary strand. Theoretically, this could suppress false positive variant calling, however this induces low library complexity, which could generate biased and incomplete sequencing data. An experimental method called IQBErepair (In vitro seQuential Base Excision repair) mimicking base excision repair process has been proposed^3^. This method remarkably improved on-target rate in WES, coverage uniformity, PCR duplication rate, however despite this improvement, more than 60% of the FFPE induced artifacts remained. Some bioinformatics tools such as FFPolish^9^, SOBDetector^10^, and Ideafix^11^ had been developed. Despite these efforts, existing methods often fall short in effectively distinguishing true somatic variants from FFPE-induced artifacts. Some filtering strategies are overly conservative and fail to adequately remove artifacts, while others mistakenly eliminate true somatic variants, particularly those with low allele frequency. As a result, FFPE-derived genomic data remain susceptible to misinterpretation, limiting their utility in clinical research.

To overcome these limitations, a more robust and targeted approach is required. Here, we present an artificial intelligence-based method specifically designed to identify and remove FFPE-induced artifacts while preserving true somatic variants. Our approach enhances the accuracy of somatic variant calling from FFPE samples, improving the reliability of genomic analyses conducted on these valuable but challenging specimens.

## Result

### Generation of paired sequencing data from fresh frozen and matched FFPE tumor samples

To investigate the characteristics of variant calls identified from FFPE specimens, we obtained 49 FFPE and their matched fresh frozen (FF) specimens from archival tumor banks of two different institutes (site A and site B). The samples from site A are from lung, colon, and rectal cancer patients operated during 2018 while those from site B are from breast and colon cancer patients operated during 2023. Whole exome sequencing (WES) was carried out with the matched 49 samples using two different exome capture kits-Agilent SureSelect V8 and Twist Exome 2.0 (hereafter referred to as Agilent kit and Twist kit, respectively) followed by variant calling using two different variant callers (MuTect2 and VarDict^12^). In addition, we used WES data from 24 cases from the public domain^6,11,13^. Whole genome sequencing (WGS) was performed on 47 matched sample pairs (two were excluded due to limited material) and somatic variant calls were generated using MuTect2 and DRAGEN^14^. The details of the analyzed specimens and the generated data are summarized in Fig. 1 and individual case characteristics are detailed in Supplementary Table 1.

**Fig. 1.**
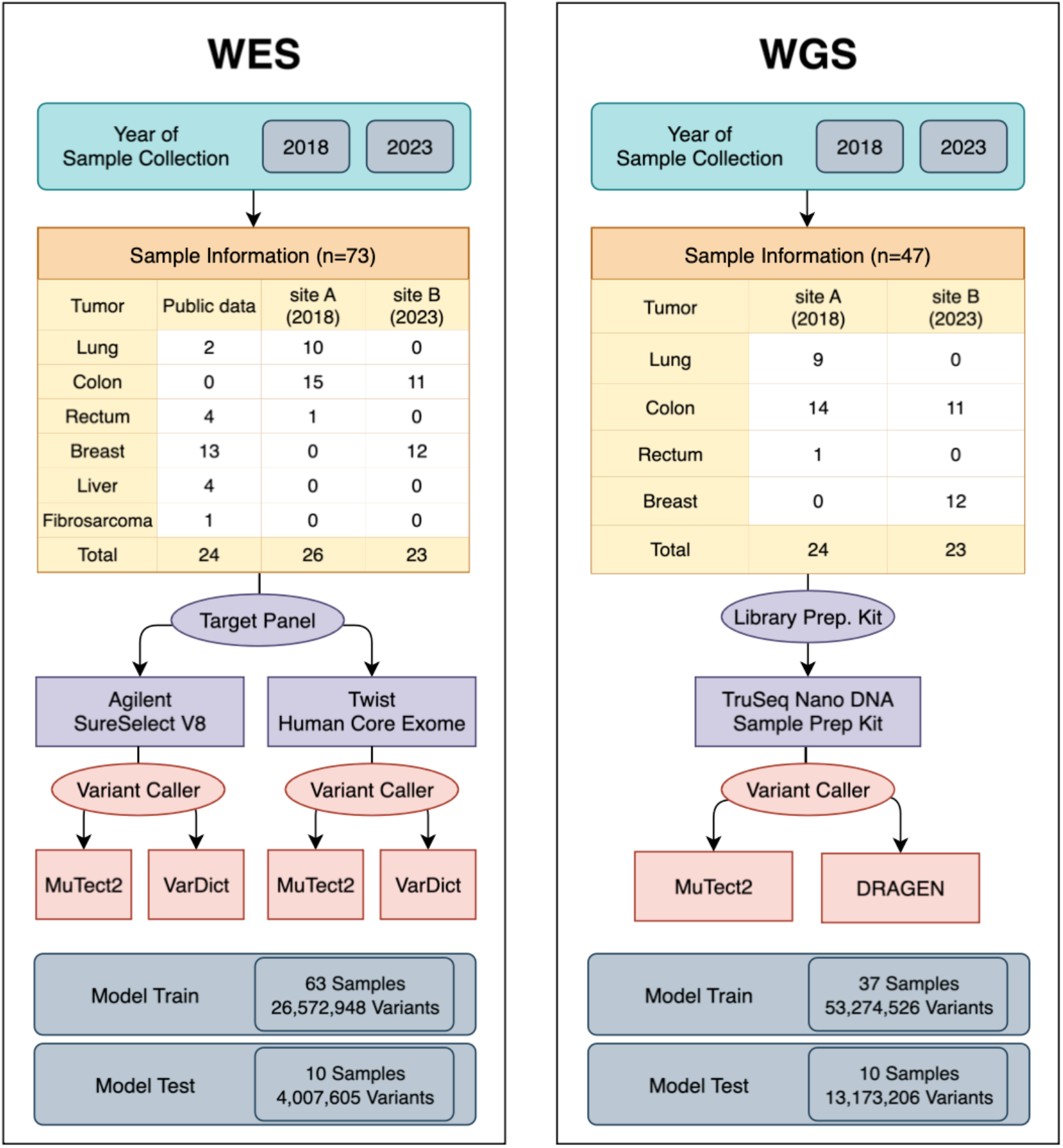
Study design and sample cohorts used in this study. The left panel summarizes the WES analysis, showing the year (site) of sample collection, the number of samples by tissue type, the exome capture kits applied, the somatic variant callers used, and the numbers of variants included for DeepOmicsFFPE-PLUS training and evaluation. The right panel presents the corresponding information for WGS analysis.

### Characteristics of the FFPE induced artifacts in the WES data

Overall, the somatic variant calls from FFPE samples differed from those in the matched fresh frozen samples (Fig. 2A, 2B). We classified the somatic variant calls into three categories; detected in FFPE only (FFPE-only), detected in both FFPE and FF (FFPE-FF both), and detected in fresh frozen-only (FF-only) (Supplementary Table 2).

**Fig. 2.**
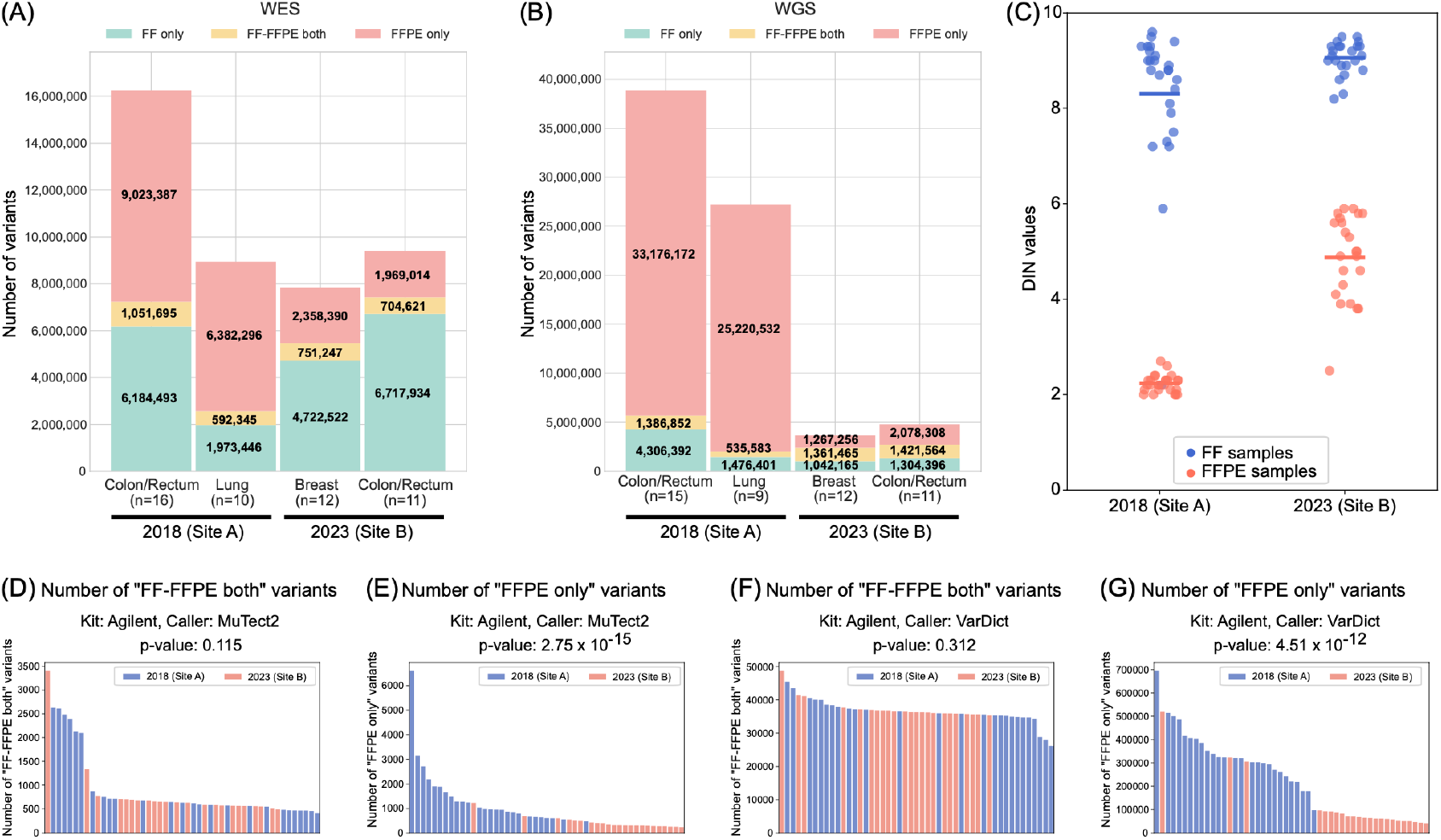
Characteristics of variant calls generated from diverse sample sources. (A, B) The number of variant calls belonging to three categories (described in the main text: ‘FF only’, ‘FF-FFPE both’, and ‘FFPE only’) from WES (A) and WGS (B) were shown. The x-axis indicates cancer types and the year (site) of sample collection. (C) Dots represent the DIN values of DNA from each sample, with blue indicating FF samples and red indicating FFPE samples. The x-axis shows the year and site of sample collection. (D-G) Samples were sorted by the number of ‘FF-FFPE both’ variants (D, F) and ‘FFPE only’ variants (E, G) and visualized with different colors according to the year (site) of sample collection. The variant calls were generated using MuTect2 (D, E) and VarDict (F, G). Statistical significance of differences across years (sites) was evaluated using negative binomial regression, with p-values indicated above each plot.

Older samples from 2018 (site A), which exhibited lower DNA integrity numbers (DINs) indicative of greater DNA damage, showed a significantly higher number of ‘FFPE-only’ variants compared to newer samples from 2023 (site B) (Fig. 2A-C). To further test this observation, we sorted each sample by its number of ‘FF-FFPE both’ variants (Fig. 2D) and ‘FFPE-only’ variants (Fig. 2E) and then visualized them in different colors according to years of collection. The number of ‘FF-FFPE both’ variants was not associated with the sample year, however, the number of ‘FFPE-only’ variants was significantly higher in the older cohort (p-value < 0.001, negative binomial regression) (Fig. 2E). These findings were consistent across variant callers (Fig. 2F, 2G) and exome capture kits (Supplementary Fig. 1A-D). Collectively, these results suggest that the ‘FFPE-only’ variants are likely artifacts related to FFPE-induced DNA damage. To further investigate this hypothesis, we looked into the strand-orientation bias of variant calls, which was known to be elevated in artifactual mutations induced by deamination or oxidation^10^. Consistent with a previous study^10^, many ‘FFPE-only’ variants exhibited higher bias scores than ‘FF-FFPE both’ variants (Supplementary Fig. 2). Since FFPE-only variants are biased toward lower variant allele frequency (VAF) (Supplementary Fig 3), they could represent subclonal mutations originating from intratumor heterogeneity. However, strong strand orientation bias of these variants suggests that they most likely represent true FFPE-induced artifacts. Based on this evidence, we defined the ‘FFPE-only’ variants as FFPE-induced artifacts that should be removed to enable accurate identification of cancer specific somatic mutations.

### In vitro DNA repair is not sufficient to eliminate FFPE-induced artifacts

It is well known that deamination of cytosine and oxidation of guanine can occur during formalin fixation, FFPE storage, and DNA extraction from FFPE blocks, resulting in artifactual C>T(C:G>T:A) and G>T(G:C>T:A) alteration, respectively^8,15^. To remedy these FFPE-induced artifacts, the treatment of uracil-DNA glycosylase and oxoguanine glycosylase has been proposed^3,5,16^. These two enzymes remove damaged bases--urical generated by cytosine deamination and 8-oxoguanine resulting from guanine oxidation--thereby preventing erroneous base pairing during DNA amplification and sequencing. To assess whether the experimental approach can sufficiently remove the artifactual variant calls from FFPE samples, we treated DNA from two FFPE samples from 2018 (site A) with a DNA repair enzyme cocktail (NEBNext FFPE DNA repair v2 module, New England Biolabs) before WES with Agilent kit. In agreement with the previous studies, C:G>T:A and G:C>T:A variants were predominantly observed among the FFPE artifacts and these artifactual alterations were specifically reduced after the treatment (Fig. 3 and Supplementary Fig. 4). However, many artifactual C:G>T:A and G:C>T:A variants, as well as other types of FFPE artifacts (in total 40 to 69% of the artifacts) still remained, suggesting that other approaches should be considered.

**Fig 3.**
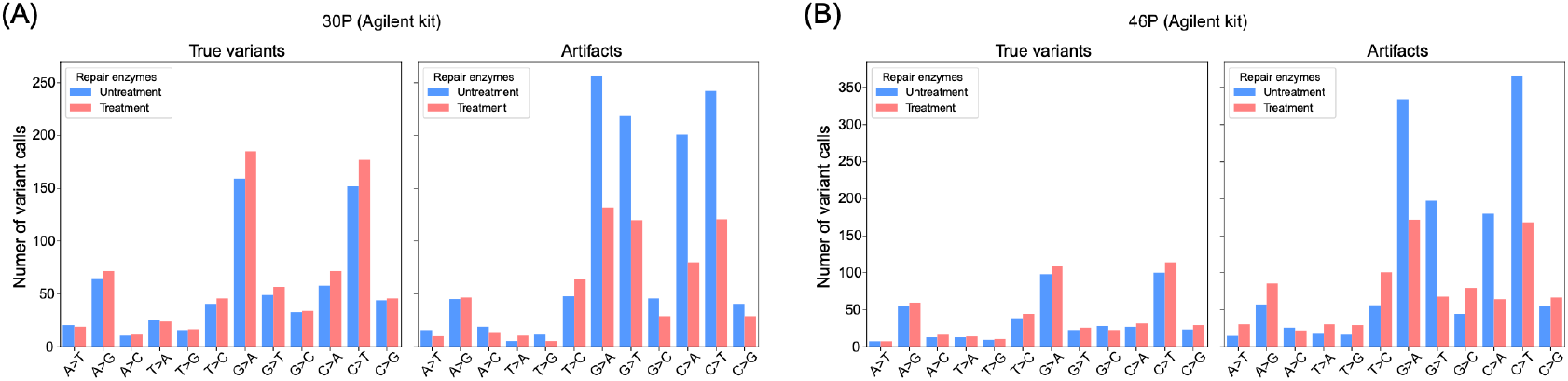
Effect of DNA repair enzyme treatment on reducing FFPE-induced artifacts in WES. Bars indicate the number of each type of single nucleotide substitution shown on the x-axis. Left and right panels show the number of true variants and artifacts, respectively, for sample 30P (A) and sample 46P (B), with or without DNA repair enzyme treatment. The Agilent exome capture kit used is indicated above each panel. The two plots within each panel (A) and (B) share the same y-axis scale.

### DeepOmicsFFPE-PLUS can improve variant calling accuracy by removing FFPE-induced artifacts in WES

We previously introduced DeepOmicsFFPE, an AI model designed to classify true variants and FFPE-induced artifacts^17^. However, a limitation of the previous version of DeepOmicsFFPE is that it only takes VCF (Variant Call Format) files generated by the MuTect2 caller. Because it relies on numeric features generated by the caller, VCF files lacking any of these features can not be used with the earlier version of DeepOmicsFFPE. Furthermore, about 12.9% of the true variants were misclassified as artifacts in the previous study. To overcome this limitation and enhance classification performance, we developed DeepOmicsFFPE-PLUS (WES) which is independent of any specific variant caller (Supplementary Fig. 5).

Based on study design (Fig. 1), using 10 samples not used for model building, we tested whether the performance of DeepOmicsFFPE-PLUS (WES) is consistent across different sample ages (2018 vs 2023), exome capture kits (Agilent vs Twist), variant callers (MuTect2 vs VarDict), as well as allele frequency. We also compared the performance of DeepOmicsFFPE-PLUS (WES) with pre-existing tools including FFPolish, SOBDetector, and the published version of DeepOmicsFFPE. Since the published version of DeepOmicsFFPE accepts only VCF files generated in the MuTect2, it was excluded from the comparison on variant calls made by VarDict caller.

We first compared the performance of each tool according to the year of sample collection (Fig. 4A, 4B). As shown in Fig. 2 and Supplementary Table 1, DNA extracted from older FFPE blocks from 2018 (site A) has lower DIN values than those from 2023 (site B), with significantly more artifacts. Results from both MuTect2 and VarDict variant calls as well as Agilent and Twist kits were combined for this analysis. A total 3,272,129 variants were detected from 2018 blocks, of which 88.2% (2,886,051) were artifacts (Fig. 4A), in contrast 735,476 variants were detected from 2023 blocks, with 484,149 artifacts (65.8%) (Fig. 4B). For 2018 and 2023 samples respectively, DeepOmicsFFPE-PLUS (WES) successfully removed 99.5% and 98.8% of the artifacts, while misclassifying only 6.2% and 4.1% of true variants, yielding a specificity of 0.995 and 0.988, and a sensitivity of 0.938 and 0.959 (Fig. 4A, 4B). These sum up to F1 scores of 0.950 and 0.968 respectively. FFPolish removed 99.8% and 99.8% of the artifacts, while misclassifying 20.7% and 20.6% of true variants as artifacts, yielding a specificity of 0.998 and 0.998, and a sensitivity of 0.793 and 0.794, and F1 scores of 0.878 and 0.883 respectively. SOBDetector was able to remove 23.4% and 34.4% of the artifacts, while misclassifying 9.5% and 9.4% of true variants as artifacts, yielding a specificity of 0.234 and 0.344, and a sensitivity of 0.905 and 0.906, and F1 scores of 0.237 and 0.572 respectively (Fig. 4A, 4B). These results demonstrated that the performance of each evaluated tool was consistent across the sample age and the severity of DNA damage. In particular, DeepOmicsFFPE-PLUS (WES) showed outstanding performance in both removing artifacts and retaining true variants regardless of the age of the samples.

**Fig 4.**
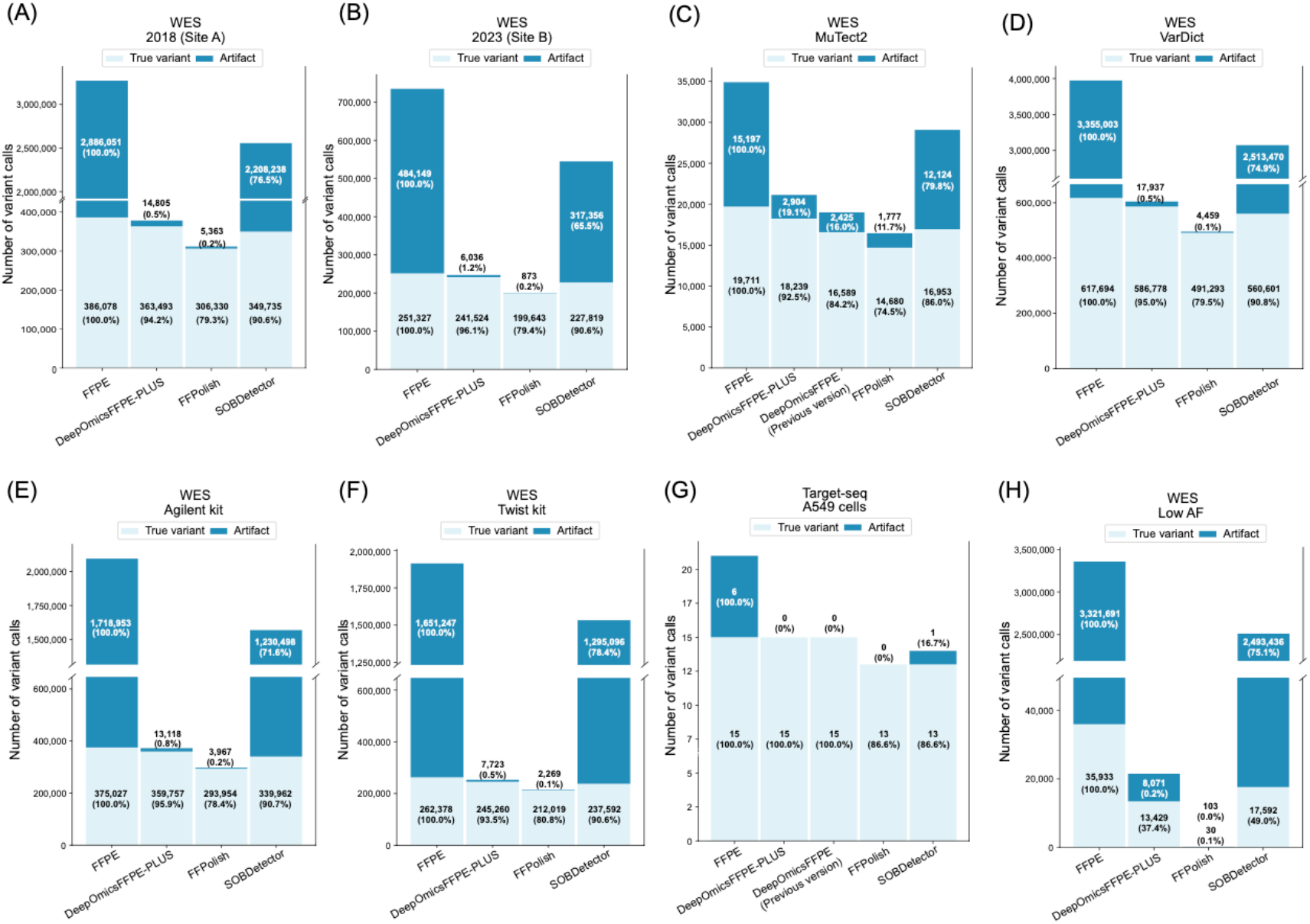
Performance evaluation of DeepOmicsFFPE-PLUS (WES) Stacked bars show FFPE-induced artifacts (dark blue) and true variants (light blue). ‘FFPE’ on the x-axis indicates unfiltered calls, while other bars represent calls after applying each tool. Performance was compared across the year (site) of sample collection (A, B), variant callers (C, D), and exome capture kits (E, F) as well as targeted sequencing of cultured A549 cells (G) and low-frequency variants (1% ≤ VAF ≤ 5%) (H).

Similarly, we evaluated the performance of the tools across two somatic variant callers (Fig. 4C, 4D) and WES kits (Fig. 4E, 4F). The earlier version of DeepOmicsFFPE removed 84.0% of artifacts (2,429 out of 15,197) and preserved 84.2% of true variants (16,589 out of 34,908) on variant calls generated using MuTect2. This result is consistent with our previous report with a different dataset^17^ (Fig. 4C). Notably, DeepOmicsFFPE-PLUS (WES) demonstrated balanced and strong performance, successfully filtering out 81.7% and 99.5% of artifacts while preserving 91.0% and 94.8% of true variants from MuTect2 and VarDict variant calls, respectively (Fig. 4C, 4D). When assessing the tool performance on variant calls from WES generated using Agilent and Twist kit (Fig. 4E, 4F), the differences were negligible. Collectively, these results indicate that DeepOmicsFFPE-PLUS (WES) consistently delivers robust and accurate performance across different WES datasets.

As previously described, FFPE-induced artifacts were defined as variants identified in an FFPE sample but not in the matched FF sample. Some variants classified as artifacts by this criterion could be subclonal true somatic variants that were not detected in the FF sample due to limitations in detection sensitivity or intratumor heterogeneity. We aimed to evaluate the classification performance of the tools using well-defined datasets generated in our previous study^17^. The targeted sequencing data were derived from paired FF and FFPE samples prepared from the same batch of cultured A549 cells. As demonstrated in the previous study, DeepOmicsFFPE outperformed FFPolish and SOBDetector on the variant calls generated by MuTect2 from the targeted sequencing data, and we observed DeepOmicsFFPE-PLUS (WES) classified all variant calls perfectly consistent with its earlier version (Fig. 4G). On variant calls generated by VarDict from the targeted sequencing data, DeepOmicsFFPE-PLUS (WES) outperformed the other tools tested in this study, achieving an F1-score of 0.964 (Supplementary Fig. 6).

### DeepOmicsFFPE-PLUS can improve variant calling accuracy of low allele frequency variants from FFPE samples

One of our goals was to ensure that DeepOmicsFFPE-PLUS (WES) could discriminate between FFPE artifacts and true somatic variants with low allele frequency. Because FFPE-induced artifacts exhibit low allele frequency (Supplementary Fig. 3), it is particularly difficult to distinguish true variants with low allele frequency from artifacts. This distinction is especially critical in contexts such as detecting drug resistance mutations following treatment, identifying neoantigen candidates in tumors with low purity or identifying emerging subclonal variants during cancer evolution. As expected, 3,321,691 out of 3,357,624 variants (98.9% of total variant calls with VAF between 0.01 and 0.05) in combined WES dataset (Agilent kit + Twist kit and MuTect2 + VarDict) were FFPE artifacts (Fig. 4H). FFPolish achieved the highest specificity (0.999), but at the cost of removing nearly all true variants (35,903 out of 35,933). In contrast, SOBDetector attained the highest sensitivity (0.435), though it suffered from the lowest specificity (0.230). DeepOmicsFFPE-PLUS offered a more balanced performance, with a specificity of 0.998 and a sensitivity of 0.337, resulting in the highest F1-score (0.439) among the tested tools. Considering all the observations, these results demonstrate DeepOmicsFFPE-PLUS (WES) can accurately distinguish true variants from FFPE-induced artifacts, even when the true variants have low allele frequencies similar to those of FFPE-artifacts.

### DeepOmicsFFPE-PLUS can improve variant calling accuracy by removing FFPE-induced artifacts in WGS

With the decreasing cost of sequencing and improved resolution for detecting structural and copy number variations, WGS is increasingly being adopted in clinical settings. To investigate whether the problematic FFPE-induced artifacts are detected in WGS data as in WES, we carried out WGS on 47 paired FF and FFPE samples (Fig. 1). Previous studies have demonstrated the superior performance of DRAGEN in variant calling^14^, even from FFPE samples^18^. However, similar to MuTect2 and VarDict used for WES, DRAGEN also called many ‘FFPE-only’ variants defined as FFPE-induced artifacts, specifically 65.8% of variant calls (6,364,525 out of 9,664,889) from FFPE samples were the artifacts (Supplementary Table 3). To examine whether increased sequencing depth improves the variant calling, we compared the variant calls from 30X and 80X WGS data across 6 samples. Although 80X sequencing led to a consistent increase in true variant detection compared to 30X, it also resulted in a substantially greater increase in artifacts (Fig. 5). This encouraged us to develop DeepOmicsFFPE-PLUS for WGS (hereafter referred to as DeepOmicsFFPE-PLUS (WGS)).

**Fig. 5.**
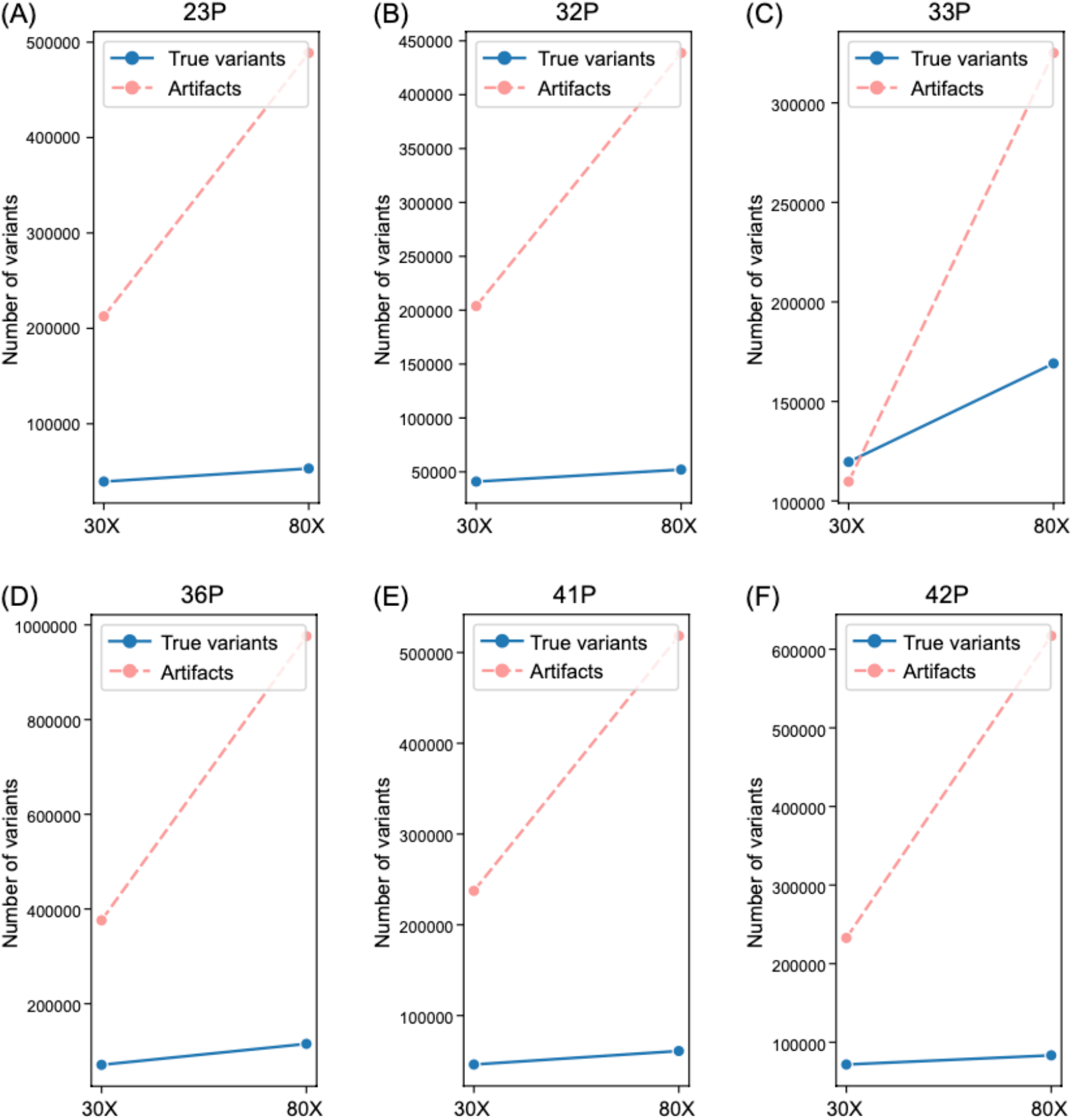
The effect of sequencing depth on variant calling accuracy. The number of true somatic variants (blue solid line) and FFPE-induced artifacts (pink dotted line) are shown for the samples indicated above each plot.

DeepOmicsFFPE-PLUS (WGS) have the same structure as DeepOmicsFFPE-PLUS (WES) (supplementary Fig. 5). We assessed the performance of DeepOmicsFFPE-PLUS (WGS) on the test datasets generated using DRAGEN (Fig. 1). With an F1-score of 0.842 on variant calls from 2018 (site A) and 0.898 on those from 2023 (site B), DeepOmicsFFPE-PLUS (WGS) exhibited consistently superior performance over FFPolish (0.553 and 0.739, respectively) and SOBDetector (0.305 and 0.874, respectively) (Fig. 6A, B). For variant calls with low variant allele frequency (0.01≦VAF≦0.05), FFPolish classified two variants as true but one of them was an artifact. In contrast, DeepOmicsFFPE-PLUS (WGS) achieved a sensitivity of 0.683, resulting in an F1-score of 0.694. SOBDetector exhibited remarkably high sensitivities of 0.864. However, it removed only 13% of artifacts (652 out of 4,985), consequently archiving an F1-score of 0.464 (Fig. 6C).

**Fig. 6.**
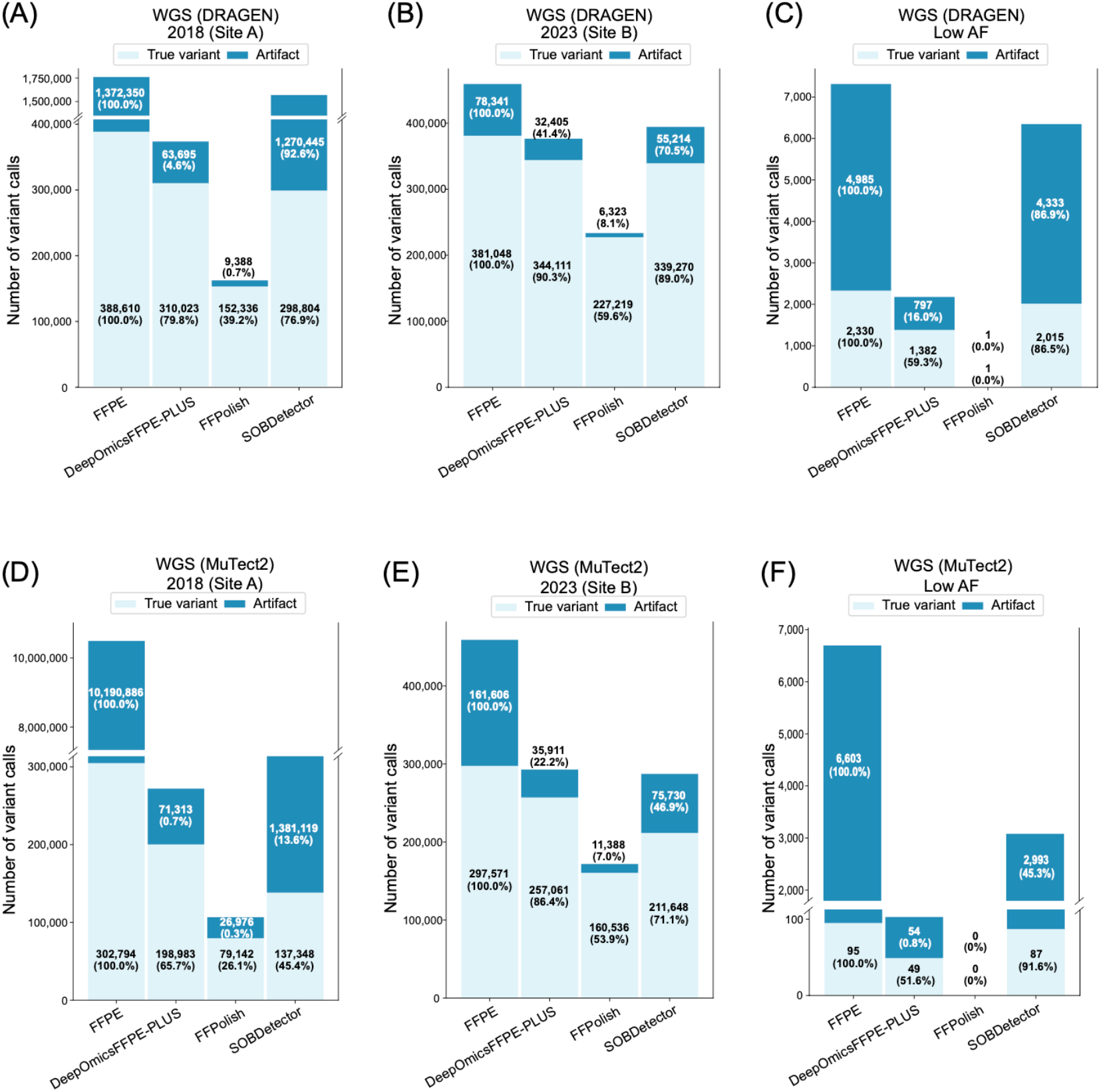
Performance evaluation of DeepOmicsFFPE-PLUS (WGS) Stacked bars represent FFPE-induced artifacts (dark blue) and true variants (light blue) as in Fig. 4. Panels A-C show results obtained using DRAGEN caller, while panels D-F use MuTect2. Panels A and B, and D and E correspond to sample collected in 2018 (site A) and 2023 (site B), respectively. Panels C and F present performance on low-frequency variants (1% ≤ VAF ≤ 5%).

Although DRAGEN provides robust variant calling, it is not as widely used as MuTect2 in many research and clinical settings. To enable broader applicability and comparability of DeepOmicsFFPE-PLUS (WGS), we additionally evaluated its performance on variant calls generated using MuTect2. Unsurprisingly, MuTect2 called a considerable number of FFPE-induced artifacts in the WGS data: specifically, 97.1% (10,190,886 out of 10,493,680) of such artifacts were detected in the 2018 samples and 35.2% (161,606 out of 459,177) in the 2023 samples. DeepOmicsFFPE-PLUS (WGS) demonstrated superior performance in distinguishing somatic variants from the artifacts compared to FFPolish and SOBDetector. It achieved an F1-score of 0.694 in the 2018 samples and 0.871 in the 2023 samples. For variant calls with low allele frequency (0.01≦VAF≦0.05) from MuTect2, FFPolish removed all variant calls, including true mutations. While SOBDetector showed the highest sensitivity (0.918), it failed to remove 45.3% of the artifacts. DeepOmicsFFPE-PLUS (WGS) demonstrated its applicability in detecting true low frequency variants by removing 99.2% of the artifacts while preserving 51.6% of true variants.

We concluded that identifying cancer-specific somatic mutations from WGS is challenging due to FFPE-induced artifacts, even with increased sequencing depth. However, DeepOmicsFFPE-PLUS (WGS) may offer a potential solution to this problem.

### DeepOmicsFFPE-PLUS offers insights into the mutational processes driving cancer development by enhancing mutational signature profiling accuracy in FFPE samples

SBS (Single Base Substitution) signatures are part of the COSMIC mutational signatures, providing insights into the etiology of cancer and the mutational processes driving tumor development^19^. For comprehensive profiling of SBS, the variant calls from WGS are typically used. Since WGS from FFPE samples harbors a substantial number of FFPE-induced artifacts, as previously demonstrated, SBS signatures of FFPE samples may differ significantly from those of their matched FF counterparts. To assess this, we profiled the SBS signatures in the FFPE and matched FF samples used to test DeepOmicsFFPE-PLUS (WGS). As expected, 7 out of 10 FFPE samples did not mirror the signatures observed in their matched FF samples. In particular, SBS signatures from FFPE samples prepared from older samples (2018) were substantially distorted (Fig. 7A, B), while those from 2023 FFPE samples closely resemble those from their matched FF samples (Fig. 7A, B). After discarding FFPE-induced artifacts (‘FFPE-only’ variants) from variant calls from each FFPE sample, the signatures were re-profiled, then we observed many suspicious signatures disappeared (Fig. 7C). A surprising thing is that the signatures associated with MSI (MicroSatellite Instability) observed in 33P FF and 36P FF samples (Fig. 7A) disappeared in their matched FFPE counterparts. It is worth noting that both 33P and 36P are derived from colon cancer, where MSI status is particularly important^20,21^. After re-profiling signatures from true variants (‘FF-FFPE both’ variants) in FFPE samples, the clinically relevant signatures were restored (Fig. 7C). This encouraged us to investigate MSI status and mutation burden of colon cancer belonging to the test set. Similar to the signatures observed in the FF samples, 33P, 36P and Colon-23 samples exhibited higher MSI scores calculated using MSIsensor2, which can accurately classify MSI status even in FFPE samples (https://github.com/niu-lab/msisensor2) (Fig. 7E). Moreover, the higher number of mutations, particularly in 33P FF, 36P FF and Colon-23 samples compared to other samples, further supports that these three samples were MSI-high (Fig. 7E). Taken together, 33P, 36P and Colon-23 samples were most likely MSI-high; however, the 33P FFPE and 36P FFPE samples did not exhibit any MSI-high related signatures and instead displayed some questionable signatures presumably related to artifacts (Fig. 7B). This implies that FFPE-induced artifacts can distort mutational signatures, potentially obscuring critical biological traits such as MSI status.

**Fig. 7.**
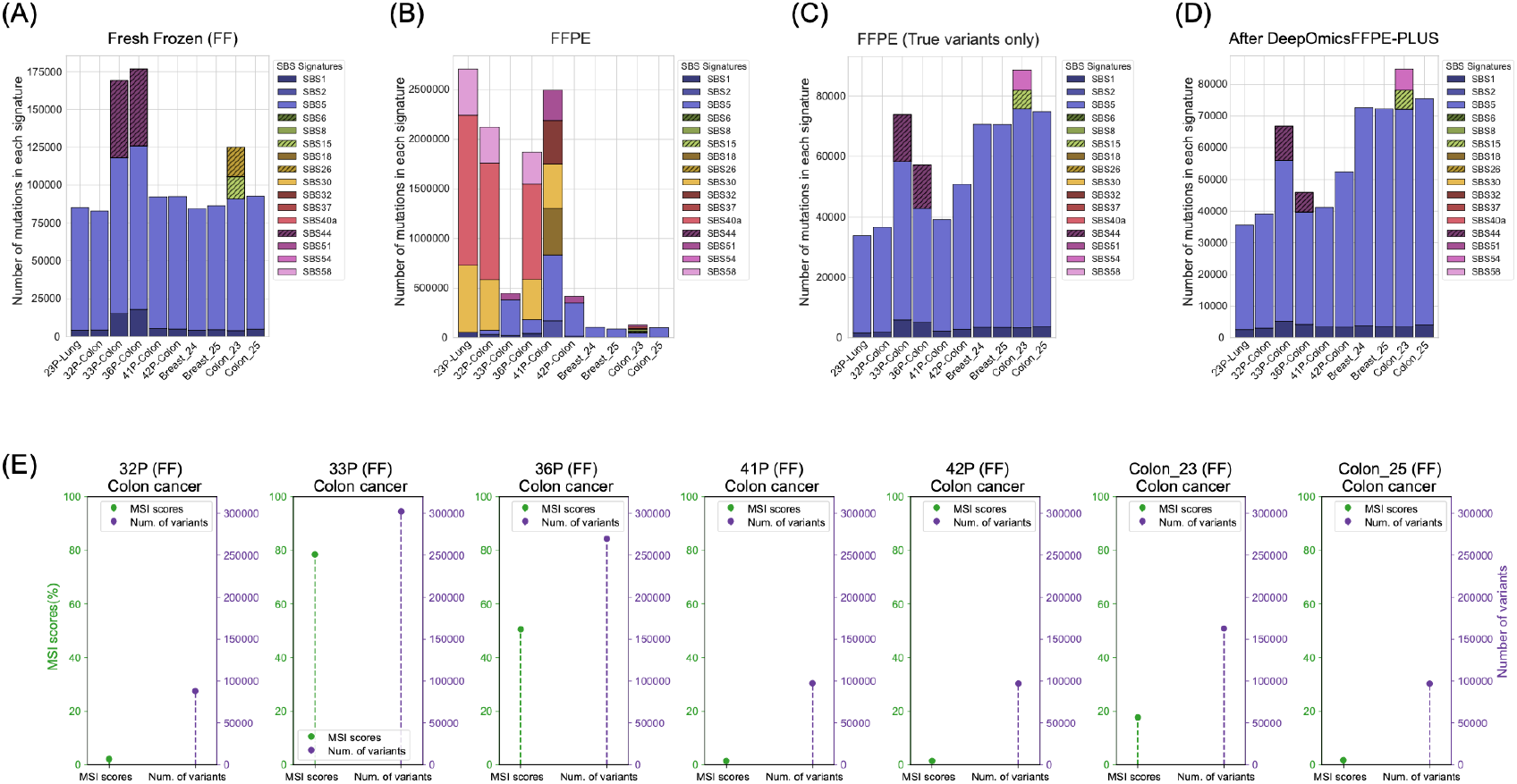
Refinement of mutational signatures from MuTect2 variant calls using DeepOmicsFFPE-PLUS. (A-D) Stacked bars show the number of variants supporting corresponding single base substitution (SBS) signatures. Samples and their tissue origins are indicated on the x-axis. MSI-associated SBS signatures (SBS6, SBS15, SBS26, and SBS44) are marked with diagonal lines. (A) SBS signatures in fresh frozen (FF) samples. (B) SBS signatures in FFPE samples before artifact removal. (C) SBS signatures in FFPE samples after removing artifacts defined as ‘FFPE only’ variants. (D) SBS signatures in FFPE samples after applying DeepOmicsFFPE-PLUS (WGS). (E) MSI scores (solid green line) and the number of variants (dotted green line) for FF colon cancer samples included in A-D. MSI scores correspond to the left y-axis, and the number of variants to the right y-axis.

To assess whether DeepOmicsFFPE-PLUS (WGS) could improve mutational signature profiles in FFPE samples, we discarded the variant calls that DeepOmicsFFPE-PLUS (WGS) classified as the artifacts and then re-profiled the mutational signatures. Surprisingly, the resulting signature profiles after applying DeepOmicsFFPE-PLUS (WGS) closely resembled those from FFPE after removing ‘FFPE-only’ variants (Fig. 7C, D). As expected, the MSI-related signatures in 33P and 36P samples were restored following DeepOmicsFFPE-PLUS (WGS) filtering (Fig. 7D). We think DeepOmicsFFPE-PLUS (WGS) can be used to refine mutational signatures by removing FFPE artifacts, particularly in the FFPE samples where DNA was severely damaged.

## Discussion

Despite the increasing adoption of WES and WGS in clinical research and diagnostics, FFPE samples continue to pose significant challenges in accurate somatic variant detection due to FFPE-induced artifacts^3,16^. This study confirms and expands upon previous findings that common variant callers, including MuTect2 and the high-performance DRAGEN, are not sufficient to resolve the confounding effect of FFPE-induced artifacts. Although DRAGEN has been known for its accuracy and computational efficiency in variant detection from high-quality samples^18^, our analysis reveals that it also suffers from high false positive rates when applied to FFPE-derived DNA, particularly in variant calls with low allele frequency.

Artifactual modifications such as cytosine deamination and guanine oxidation are well-known contributors to FFPE-induced artifacts, resulting in C:G>T:A and G:C>T:A substitutions^3^. However, other sources of artifact, such as hairpin-like secondary structures or single-strand overhangs, can also arise during FFPE DNA extraction and NGS library preparation from damaged DNA, producing various types of false positive variant calls^22^. This mechanistic insight may explain why DNA repair enzyme treatment exhibited only a partial effect on removing FFPE-induced artifacts (Fig. 3). Moreover, we observed that increasing sequencing depth from 30X to 80X led to a consistent increase in the number of true variant calls; however, this was accompanied by an even more pronounced rise in artifact calls. This disproportionate increase indicates that higher depth alone is not an adequate solution and may, in fact, amplify the misleading influence of the artifacts (Fig. 5). These findings underscore the critical need for computational correction mechanisms that go beyond enzymatic correction and simple depth scaling.

We demonstrated that DeepOmicsFFPE-PLUS significantly outperformed existing tools such as FFPolish and SOBDetector across various settings, including WES, WGS, FFPE blocks of different ages, exome capture kits, somatic variant callers, and low allele frequency variant calls. Moreover, we showed that DeepOmicsFFPE-PLUS substantially improves the fidelity of downstream analysis such as mutational signature profiling. By removing FFPE-induced artifacts from WGS variant calls, we recovered SBS profiles that closely mirrored those from matched FF samples, a crucial step for accurate biological interpretation. Notably, in cases of some colon cancers (33P and 36P), DeepOmicsFFPE-PLUS not only eliminated spurious SBS signatures but also restored biologically relevant signatures that were initially obscured (Fig. 7). These findings suggest that DeepOmicsFFPE-PLUS can facilitate the interpretation of disease-related etiologies and mutational processes underlying cancer evolution by effectively removing FFPE-induced artifacts. It is also important to note that mutational signature profiling has become an essential tool in understanding cancer etiology and guiding treatment strategies, particularly in cancers of unknown primary (CUP)^23,24^. Thus DeepOmicsFFPE-PLUS may serve as a critical enabler of signature-driven clinical applications, including CUP diagnostics and personalized treatment planning.

In summary, DeepOmicsFFPE-PLUS has the potential to facilitate accurate somatic variant detection across a wide range of applications, including studies of cancer evolution, identification of neoantigens for cancer vaccine development, detection of resistance mutations following therapeutic interventions, and characterization of hotspot mutations in low-purity tumors. Its robustness also makes it suitable for retrospective analyses using archived FFPE samples, broadening its utility in both clinical and translational cancer research.

## Methods

### Biomaterials

This study was approved by the Institutional Review Boards of Seoul St. Mary’s Hospital and Konkuk University Medical Center. A total of 26 FFPE blocks and 23 FFPE blocks as well as corresponding paired FF tissues were obtained from the Biobank of Seoul St. Mary’s Hospital, The Catholic University of Korea, and Konkuk University Medical Center, respectively. These samples included 10 lung cancers, 26 colon cancers, 1 rectal cancer, and 12 breast cancers.

DNA was extracted from the FFPE blocks using ReliaPrep FFPE gDNA Miniprep System (Promega) and from FF tissues using QIAamp DNA mini kit (QIAgen) in accordance with the manufacturer’s instructions. DIN values of extracted DNA were calculated using the TapeStation system (Agilent). For DNA repair enzyme treatment, 0.2 ug of DNA was treated with NEBNext FFPE DNA repair module v2 (New England Biolabs) according to the manufacturer’s instructions.

### Whole exome sequencing

A total 49 FFPE samples and their matched FF samples were subjected to whole exome sequencing (WES) (Supplementary Table 1). Exome capture and library preparation were performed using SureSelect human all exome V8 kit (Agilent) and Exome 2.0 kit (Twist), following the manufacturer’s instructions. Sequencing was carried out on a NovaSeq X Plus sequencer (Illumina), generating an average of 30 gigabases per sample.

Reads were aligned to the human reference genome (hg38 assembly) using BWA (Burrows-Wheeler Aligner) and potential PCR duplicates were removed with Picard. Base Quality Score Recalibration (BQSR) was performed using the GATK toolkit, followed by variant calling using MuTect2 (Broad Institute) and VarDict (AstraZeneca)^12^. Only variants marked as ‘PASS’ in the FILTER column of VCF (Variant Call Format) files were considered for further analysis using bcftools^25^. The same variant detected by different kits or variant callers from the same sample was treated as independent data and used for training and testing.

### Whole genome sequencing

Due to the limited quantities of 22P and 47P from site A, whole genome sequencing (WGS) was performed on 47 FFPE samples and their matched FF counterparts (Supplementary Table 1). Library preparation for WGS was performed using the TruSeq DNA Nano kit (Illumina), following the manufacturer’s instructions. The libraries were then sequenced on a NovaSeq X Plus sequencer (Illumina), generating an average of 100 gigabases per sample for 30X sequencing depth and 260 gigabases for 80X sequencing depth.

Read alignment and variant calling were carried out using DRAGEN (Illumina) on the Illumina Connected Analytics (ICA) platform. The resulting files were downloaded, and only variants marked as ‘PASS’ in the FILTER column and annotated with ‘GermlineStatus=Somatic’ in the INFO column were retained for further analysis. The same WGS data were also analyzed using the GATK toolkit with MuTect2 for variant calling, and variants marked as ‘PASS’ tag in the FILTER column in VCFs were used in this study. The corresponding variants were extracted using bcftools^25^ from VCFs.

### Developing DeepOmicsFFPE-PLUS and performance evaluation

DeepOmicsFFPE-PLUS consists of an encoder inspired by the Transformer model^26^ and four fully connected layers. The encoder comprises four self-attention blocks, skip connections, normalization layers, and feed-forward layers. This encoder takes an 11bp sequence as input and returns a 1,024-dimensional vector. Variant types such as single nucleotide variations, and short insertions and deletions are embedded into a 4-dimensional vector. The concatenated vector, comprising the sequence embedding, the variant type embedding, and numeric features of a variant call, is passed through the fully connected layers. The first three layers each use a rectified linear unit (ReLU) activation function^27^, while the last layer uses a sigmoid activation function. Two batch normalization layers are applied between the first and second layer, and between the third and final layer^28^. Binary Cross Entropy Loss (BCLoss) was used to calculate the loss, and the Adam optimizer was used to update the network weights^29^. The model was implemented with PyTorch (version 1.11.10)^30^. DeepOmicsFFPE-PLUS is available at [github address]

To train DeepOmicsFFPE-PLUS (WES), we used 20 of 26 samples from site A, 19 of 23 samples from site B, and 24 publicly available WES datasets. The remaining 6 and 4 samples, respectively, were used to test DeepOmicsFFPE-PLUS (WES) after training it. For DeepOmicsFFPE-PLUS (WGS), the exact same samples as used for the WES model were employed, except two samples, as described earlier. To train each model, eight NVIDIA A100 graphics processing units (GPUs) were used.

The same datasets were used to evaluate the performance of DeepOmicsFFPE-PLUS, DeepOmicsFFPE (the earlier version), FFPolish^9^, and SOBDetetor^10^. Since the earlier version of DeepOmicsFFPE is only compatible with the output from MuTect2 caller, it was included only in the comparisons using the variant calls generated by MuTect2. Accuracy, specificity, sensitivity, precision, and F1-score were calculated as follows:

Accuracy = (TP + TN) / total number of variant calls

Specificity = TN / (TN + FP)

Sensitivity = TP / (TP + FN)

Precision = TP / (FP + TP)

F1-score = 2 x sensitivity x precision / (sensitivity + precision)

where TP, TN, FP, and FN represent the number of true positives, true negatives, false positives, and false negatives, respectively.

### Exploratory data analysis and visualization

SOB scores of individual variant calls were computed as previously described using an in-house script implemented in Python. VAFs were extracted from the VCF files. All plots in this manuscript were generated using Python packages matplotlib (version 3.5.2) and seaborn (version 0.11.2).

### Microsatellite analysis

6 colon cancer WGS data belonging to the test dataset were subjected to microsatellite analysis using MSIsensor2 (https://github.com/niu-lab/msisensor2, version 0.1) with default settings.

### Mutational signature profiling

To characterize mutational signatures of 10 WGS data belonging to the test dataset, SigProfilerAssignment (https://github.com/alexandrovlab/SigProfilerAssignment, version 0.1.7) was used with COSMIC version 3.4.

## Author contributions

D.-H.H, S.-E.H and S.P. conceived and designed the study. D.-H.H and S.P wrote the manuscript. W.S.K, A.L. provided the resources for the study and did data curation. D.-H.H and I.K. were developed the core algorithm of DeepOmicsFFPE-PLUS, assessed the performance of it and visualized all data used in the manuscript. I.K. and J.P. conducted variant calling from WES and WGS data. J.L., E.N., M.K., J.P., J.H., M.P., and H.C conducted WES/WGS. J.S. and Y.Y were developed the user interface of DeepOmicsFFPE-PLUS.

## Acknowledgements

This research was supported by a grant of Korean ARPA-H Project through the Korea Health Industry Development Institute (KHIDI), funded by the Ministry of Health & Welfare, Republic of Korea (grant number: RS-2025-25402727).

