## Supplementary Table 1 for "Accurate somatic variant detection from formalin fixed, paraffin embedded tissue (FFPE) derived WES and WGS by DeepOmicsFFPE-PLUS, a sequence context-based transformer"

Supplementary Table 1. Details of the specimens used in this study.

| FFPE sample | DIN from FFPE sample | Matched FF sample | DIN from FF sample | Tumor | Year of sample collection | Site | Train or Test for DeepOmicsFFPE-PLUS |
| --- | --- | --- | --- | --- | --- | --- | --- |
| 22P | 2 | 180212_001_22P | 2.2 | Lung | 2018 | A | Train (WES) / Not used (WGS) |
| 23P | 2.1 | 180213_013_23P | 9.3 | Lung | 2018 | A | Test |
| 24P | 2 | 180626_023_24P | 9.3 | Lung | 2018 | A | Train |
| 25P | 2.4 | 180906_001_25P | 9 | Lung | 2018 | A | Train |
| 26P | 2.1 | 180102_002_26P | 9.5 | Lung | 2018 | A | Train |
| 27P | 2 | 180326_010_27P | 9.4 | Lung | 2018 | A | Train |
| 28P | 2.1 | 180515_011_28P | 8.7 | Lung | 2018 | A | Train |
| 29P | 2 | 180611_020_29P | 7.3 | Lung | 2018 | A | Train |
| 30P | 2.2 | 180726_016_30P | 5.9 | Lung | 2018 | A | Train |
| 31P | 2.3 | 180921_020_31P | 9.3 | Lung | 2018 | A | Train |
| 32P | 2.1 | 180627_001_32P | 8.8 | Colon | 2018 | A | Test |
| 33P | 2.7 | 180725_007_33P | 8.1 | Colon | 2018 | A | Test |
| 34P | 2.2 | 180214_010_34P | 7.5 | Colon | 2018 | A | Train |
| 35P | 2 | 171002_013_35P | 9.6 | Colon | 2018 | A | Train |
| 36P | 2.2 | 170908_010_36P | 8.6 | Colon | 2018 | A | Test |
| 37P | 2.2 | 170720_008_37P | 8.4 | Colon | 2018 | A | Train |
| 38P | 2.2 | 170711_011_38P | 7.9 | Colon | 2018 | A | Train |
| 39P | 2.3 | 180723_016_39P | 9 | Colon | 2018 | A | Train |
| 40P | 2.4 | 180720_011_40P | 8.9 | Colon | 2018 | A | Train |
| 41P | 2.3 | 180719_001_41P | 9 | Colon | 2018 | A | Test |
| 42P | 2.6 | 180807_016_42P | 7.2 | Colon | 2018 | A | Test |
| 43P | 2.3 | 180625_021_43P | 8.8 | Colon | 2018 | A | Train |
| 44P | 2.3 | 180607_022_44P | 9.2 | Colon | 2018 | A | Train |
| 45P | 2.4 | 180621_012_45P | 9.1 | Rectum | 2018 | A | Train |
| 46P | 2.3 | 180705_017_46P | 8.8 | Colon | 2018 | A | Train |
| 47P | 2.3 | 181002_023_47P | 7.2 | Colon | 2018 | A | Train (WES) / Not used (WGS) |
| Breast_13_FFPE | 5.7 | Breast_13_FROZ | 9.3 | Breast | 2023 | B | Train |
| Breast_14_FFPE | 4.9 | Breast_14_FROZ | 9.2 | Breast | 2023 | B | Train |
| Breast_15_FFPE | 5.8 | Breast_15_FROZ | 9.1 | Breast | 2023 | B | Train |
| Breast_16_FFPE | 5.6 | Breast_16_FROZ | 9 | Breast | 2023 | B | Train |
| Breast_18_FFPE | 5.3 | Breast_18_FROZ | 9.5 | Breast | 2023 | B | Train |
| Breast_19_FFPE | 5.8 | Breast_19_FROZ | 9.3 | Breast | 2023 | B | Train |
| Breast_20_FFPE | 4.6 | Breast_20_FROZ | 9.3 | Breast | 2023 | B | Train |
| Breast_21_FFPE | 3.8 | Breast_21_FROZ | 9.3 | Breast | 2023 | B | Train |
| Breast_22_FFPE | 5.4 | Breast_22_FROZ | 9.2 | Breast | 2023 | B | Train |
| Breast_23_FFPE | 5 | Breast_23_FROZ | 9.3 | Breast | 2023 | B | Train |
| Breast_24_FFPE | 5.6 | Breast_24_FROZ | 9.4 | Breast | 2023 | B | Test |
| Breast_25_FFPE | 5.9 | Breast_25_FROZ | 9.5 | Breast | 2023 | B | Test |
| Colon_14_FFPE | 4.3 | Colon_14_FROZ | 8.8 | Colon | 2023 | B | Train |
| Colon_15_FFPE | 3.8 | Colon_15_FROZ | 9.4 | Colon | 2023 | B | Train |
| Colon_16_FFPE | 5 | Colon_16_FROZ | 9.1 | Colon | 2023 | B | Train |
| Colon_17_FFPE | 4.9 | Colon_17_FROZ | 8.9 | Colon | 2023 | B | Train |
| Colon_18_FFPE | 4.6 | Colon_18_FROZ | 8.3 | Colon | 2023 | B | Train |
| Colon_19_FFPE | 2.5 | Colon_19_FROZ | 9 | Colon | 2023 | B | Train |
| Colon_20_FFPE | 3.9 | Colon_20_FROZ | 8.6 | Colon | 2023 | B | Train |
| Colon_21_FFPE | 3.9 | Colon_21_FROZ | 9 | Colon | 2023 | B | Train |
| Colon_22_FFPE | 4.1 | Colon_22_FROZ | 8.7 | Colon | 2023 | B | Train |
| Colon_23_FFPE | 5.9 | Colon_23_FROZ | 8.9 | Colon | 2023 | B | Test |
| Colon_25_FFPE | 5.8 | Colon_25_FROZ | 8.2 | Colon | 2023 | B | Test |

Supplementary Table 2. Details of variants identified by WES

| Train/Test | Year (Site) | Number of samples | Target panel | Variant caller | Number of variants | | | |
| --- | --- | --- | --- | --- | --- | --- | --- | --- |
|  |  |  |  |  | Total | FF only | FF-FFPE both | FFPE only |
| Train | 2018 (site A) | 20 | Agilent | MuTect2 | 63,773 | 13,390 | 18,789 | 31,594 |
|  |  |  |  | VarDict | 12,095,019 | 4,208,952 | 724,138 | 7,161,929 |
|  |  |  | Twist | MuTect2 | 82,221 | 9,369 | 14,795 | 58,057 |
|  |  |  |  | VarDict | 8,055,324 | 2,287,032 | 501,060 | 5,267,232 |
|  | 2023 (site B) | 19 | Agilent | MuTect2 | 29,015 | 6,584 | 14,640 | 7,791 |
|  |  |  |  | VarDict | 8,403,199 | 5,436,630 | 713,296 | 2,253,273 |
|  |  |  | Twist | MuTect2 | 23,150 | 6,483 | 10,892 | 5,775 |
|  |  |  |  | VarDict | 7,438,312 | 5,396,183 | 466,287 | 1,575,842 |
|  | Public data | 24 | - | MuTect2 | 101,788 | 11,121 | 19,431 | 71,236 |
|  |  |  |  | VarDict | 9,435,118 | 1,867,506 | 726,718 | 6,840,894 |
| Test | 2018 (site A) | 6 | Agilent | MuTect2 | 16,143 | 3,585 | 7,290 | 5,268 |
|  |  |  |  | VarDict | 2,520,157 | 799,798 | 218,923 | 1,501,436 |
|  |  |  | Twist | MuTect2 | 16,941 | 2,629 | 5,514 | 8,798 |
|  |  |  |  | VarDict | 2,358,084 | 833,184 | 153,531 | 1,371,369 |
|  | 2023 (site B) | 4 | Agilent | MuTect2 | 5,840 | 1,169 | 3,141 | 1,530 |
|  |  |  |  | VarDict | 551,722 | 195,330 | 144,985 | 211,407 |
|  |  |  | Twist | MuTect2 | 4,376 | 1,009 | 2,372 | 995 |
|  |  |  |  | VarDict | 768,114 | 397,068 | 100,255 | 270,791 |
| Total | | 73 | - | - | 51,968,296 | 21,477,022 | 3,846,057 | 26,645,217 |

Supplementary Table 3. Details of variants identified by WGS

| Train/Test | Year (Site) | Number of samples | Variant caller | Number of variants | | | |
| --- | --- | --- | --- | --- | --- | --- | --- |
|  |  |  |  | Total | FF only | FF-FFPE both | FFPE only |
| Train | 2018 (site A) | 18 | MuTect2 | 44,619,655 | 1,696,754 | 735,016 | 42,187,885 |
|  |  |  | DRAGEN | 7,024,630 | 1,883,032 | 842,930 | 4,298,668 |
|  | 2023 (site B) | 19 | MuTect2 | 3,548,419 | 641,334 | 1,300,618 | 1,606,467 |
|  |  |  | DRAGEN | 2,712,158 | 409,216 | 1,687,776 | 615,166 |
| Test | 2018 (site A) | 6 | MuTect2 | 11,134,104 | 640,424 | 302,794 | 10,190,886 |
|  |  |  | DRAGEN | 2,499,002 | 738,042 | 388,610 | 1,372,350 |
|  | 2023 (site B) | 4 | MuTect2 | 600,735 | 141,558 | 297,571 | 161,606 |
|  |  |  | DRAGEN | 546,144 | 86,755 | 381,048 | 78,341 |
| Total | - | 47 | - | 72,684,847 | 6,237,115 | 5,936,363 | 60,511,369 |
