## Supplementary figures and legend for "Accurate somatic variant detection from formalin fixed, paraffin embedded tissue (FFPE) derived WES and WGS by DeepOmicsFFPE-PLUS, a sequence context-based transformer"

**Supplementary Figures and legends**

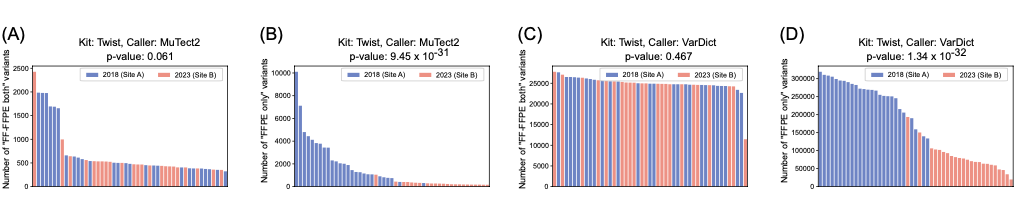

**Supplementary Fig. 1. Characteristics of variant calls generated from diverse sample sources.**

Samples were sorted by the number of “FF-FFPE both” variants (A, C) and “FFPE only” variants (B, D) and visualized with different colors according to the year (site) of sample collection. The variant calls were generated using MuTect2 (A, B) and VarDict (C, D). Statistical significance of differences across years (sites) was evaluated using negative binomial regression, with p-values indicated above each plot.

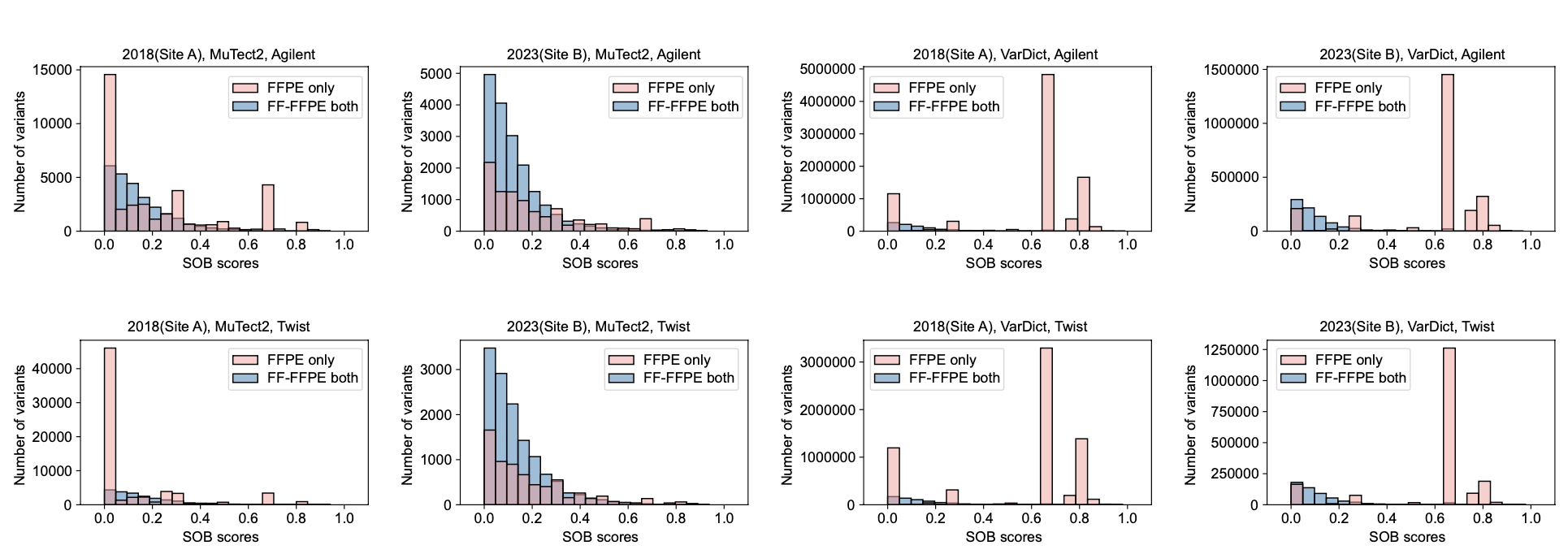

**Supplementary Fig. 2. Strand-orientation bias (SOB) scores of “FFPE only” and “FF-FFPE both” variants.**

The SOB score and corresponding the number of variants are shown on the x- and the y-axis, respectively. The upper panels present the variant calls from WES data generated using the Agilent exome capture kit, while the lower panels show those obtained with the Twist kit. The year (site) of sample collection and variant callers used are indicated above each plot.

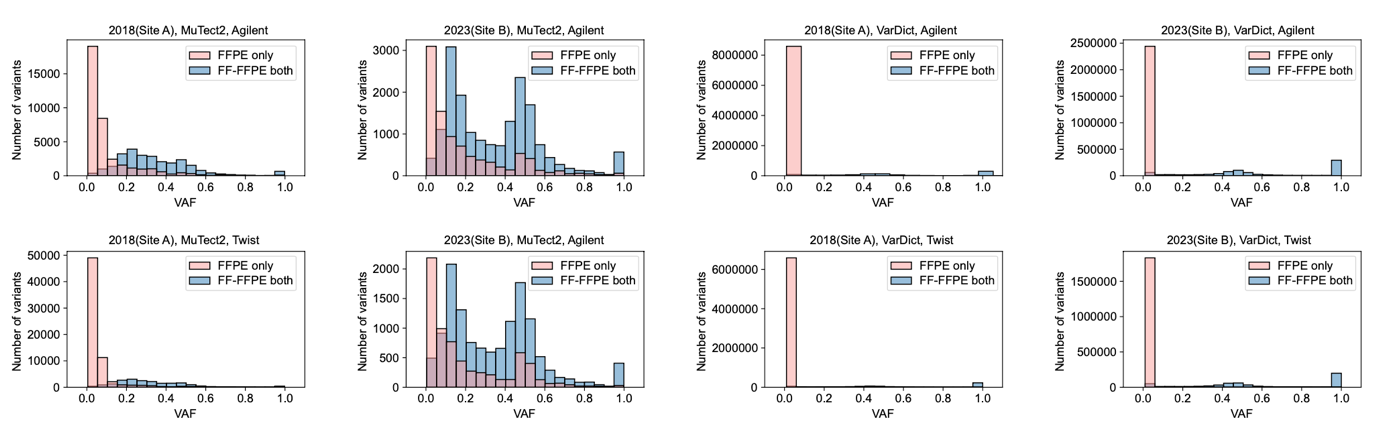

**Supplementary Fig. 3. Variant allele frequencies (VAF) of “FFPE only” and “FF-FFPE both” variants.**

The VAF and corresponding the number of variants are shown on the x- and the y-axis, respectively. The upper panels present the variant calls from WES data generated using the Agilent exome capture kit, while the lower panels show those obtained with the Twist kit. The year (site) of sample collection and variant callers used are indicated above each plot.

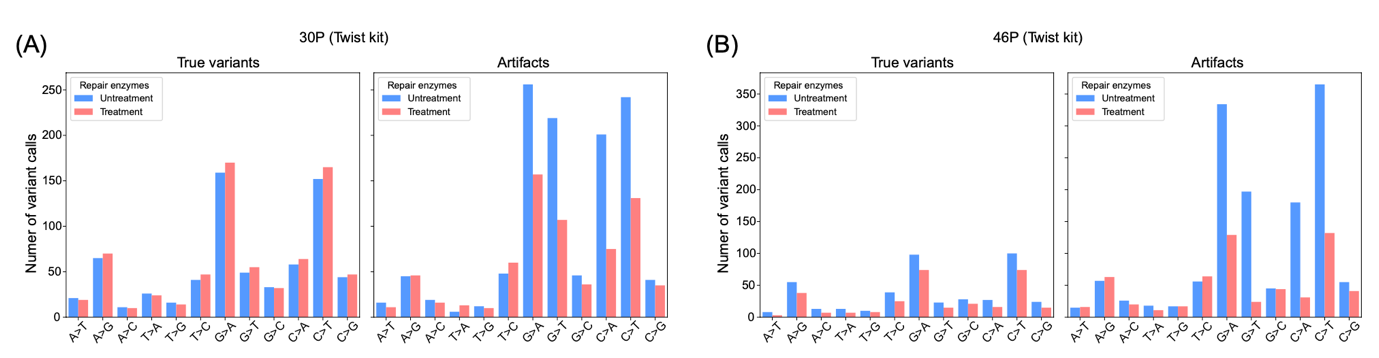

**Supplementary Fig. 4. Effect of DNA repair enzyme treatment on reducing FFPE-induced artifacts in WES.**

Bars indicate the number of each type of single nucleotide substitution shown on the x-axis. Left and right panels show the number of true variants and artifacts, respectively, for sample 30P (A) and sample 46P (B), with or without DNA repair enzyme treatment. The Twist exome capture kit used is indicated above each panel. The two plots within each panel (A) and (B) share the same y-axis scale.

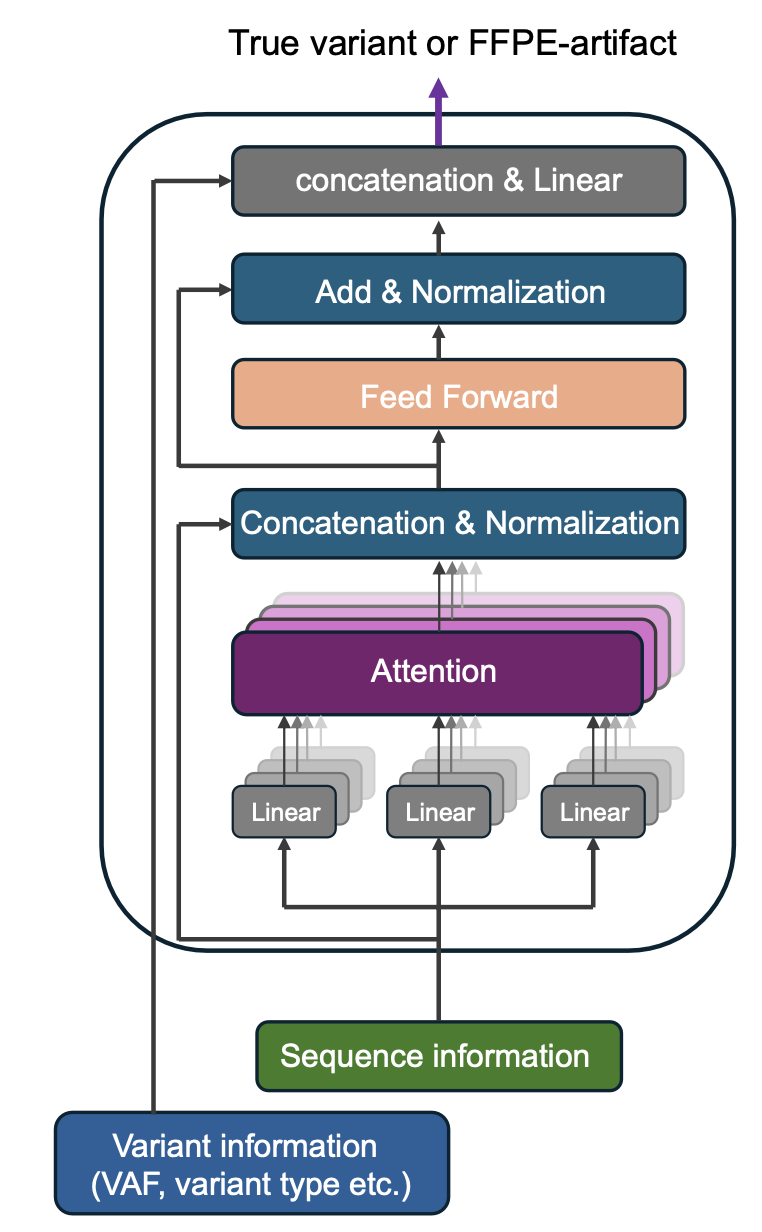

**Supplementary Fig. 5. Schematic illustration of the DeepOmicsFFPE-PLUS architecture.**

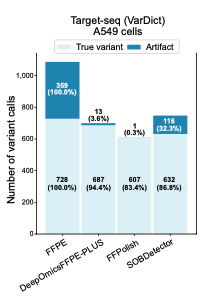

**Supplementary Fig. 6. Performance evaluation of DeepOmicsFFPE-PLUS (WES)**

Stacked bars show FFPE-induced artifacts (dark blue) and true variants (light blue). ‘FFPE’ on the x-axis indicates unfiltered calls, while other bars represent calls after applying each tool. Performance was compared on variant calls generated by VarDict from targeted sequencing of cultured A549 cells.
